# Attention-related modulation of caudate neurons depends on superior colliculus activity

**DOI:** 10.1101/843235

**Authors:** James P. Herman, Fabrice Arcizet, Richard J. Krauzlis

## Abstract

Recent work has implicated the basal ganglia in visual perception and attention, in addition to their traditional role in motor control. The basal ganglia, especially the caudate nucleus “head” (CDh) of the striatum, receive indirect anatomical connections from the superior colliculus, a midbrain structure that is known to play a crucial role in the control of visual attention. To test the possible functional relationship between these subcortical structures, we recorded CDh neuronal activity before and during unilateral SC inactivation in a spatial attention task. SC inactivation significantly altered the attention-related modulation of CDh neurons and strongly impaired the classification of task epochs based on CDh activity. Only inactivation of the same-side of SC as recorded CDh neurons, not the opposite-side, had these effects. These results demonstrate a novel interaction between SC activity and attention-related visual processing in the basal ganglia.

## Introduction

Covert visual attention is the ability of primates to selectively make use of some visual inputs while ignoring the rest, without moving the eyes. It confers exquisite behavioral and cognitive flexibility. For example, a male macaque of lower social rank can simultaneously try to appease a higher-ranking male by avoiding direct eye contact and monitor the higher-ranking male’s behavior for signs of overt aggression. In laboratory tests, covert attention is reliably associated with changes in the speed and accuracy of behavioral reports (Carrasco, 2011). The neural mechanisms for covert visual attention in primates include the modulation of neuronal activity in visual cortical areas that represent stimulus features (Reynolds and Chelazzi, 2004; Treue, 2001), as well as areas of the frontal and parietal cortex that regulate what is attended (Bisley and Goldberg, 2010; Moore and Zirnsak, 2017). More recently, it has been recognized that in addition to these cortical mechanisms, the control of covert attention also includes subcortical brain regions, consistent with the idea that covert attention in primates evolved from older subcortical circuits responsible for overt orienting movements triggered by stimulus events (Krauzlis et al., 2018).

One of the most important subcortical structures for the control of covert visual attention in primates is the superior colliculus (SC), located in the midbrain. When SC neuronal activity is perturbed by inactivation or microstimulation, performance in attention tasks is reliably altered in a spatially specific manner, even during covert tasks (Bogadhi et al., 2019; Bollimunta et al., 2018; Cavanaugh and Wurtz, 2004; Herman et al., 2018; Lovejoy and Krauzlis, 2010; Müller et al., 2005; Zénon and Krauzlis, 2012). During SC inactivation, improvements in perceptual sensitivity made possible by spatial cues are abolished (Lovejoy and Krauzlis, 2017). Despite SC inactivation preventing spatial cues from conferring a perceptual advantage, cue-related modulation of neuronal activity in extrastriate visual cortex remains robust during inactivation (Zénon and Krauzlis, 2012). Specifically, Zénon & Krauzlis (2012) found that in a covert motion-change detection task, neurons in visual areas MT and MST showed the same cue-related modulation during SC inactivation as they had before. Because behavior in this task depends on signals arising from MT / MST, these results suggest that SC inactivation impairs behavior by altering the use of cortical sensory signals in another brain area. One candidate site, based on the convergence of signals from visual cortex and the SC, is the striatum of the basal ganglia (Redgrave et al., 2010; Saint-Cyr et al., 1990), in particular, the “head” of the caudate nucleus (CDh).

The caudate is a primary input nucleus of the basal ganglia with distinct territories that have been implicated in value-based decision making, perceptual choices and selective attention. The caudate is divided into the “head” (CDh) at the anterior end, followed by the “body”, “genu”, and “tail” (CDt). In keeping with the parallel functional circuit architecture of the basal ganglia (Alexander et al., 1986), these territories appear to have distinct functional roles. For example, inactivating CDh neurons impairs choice among visual stimuli that are flexibly associated with high or low reward but does not impair choice of stimuli with fixed reward association, whereas CDt inactivation impairs only choices with fixed reward association stimuli and not flexible reward (Kim and Hikosaka, 2013). Microstimulation of CDh and anterior caudate body neurons spatially biases perceptual decisions and alters decision times with random-dot-motion stimuli, consistent with a role for CDh neurons in linking cortical visual signals, perceptual choice, and spatial selection (Ding and Gold, 2012). Finally, during an attention task requiring monkeys to perform covert perceptual judgments, CDh and body neurons are strongly modulated by the location of a spatial cue, response choice, or both (Arcizet and Krauzlis, 2018). These results suggest that CDh neuronal activity is driven by a combination of sensory signals, information about behavioral relevance, spatial location and response choice.

We hypothesized that activity in the intermediate and deep layers of primate SC contributes to cue-related modulation of neuronal activity in the caudate nucleus. To test this idea, we recorded the activity of populations of CDh neurons with a pair of linear electrode arrays while monkeys performed a covert attention task - both before and during unilateral inactivation of the SC. Our results demonstrate that inactivation of SC on the “same-side” of the brain as recorded CDh neurons changes how those neurons represent stimulus relevance while monkeys are performing covert perceptual judgments. We find that same-side SC inactivation: (1) causes clear shifts in the cue-side preferences of CDh neurons; (2) disrupts the ability of a classifier to uniquely identify distinct task-epochs on the basis of CDh neuronal activity; and (3) alters the structure of correlations in CDh neuron populations, consistent with reducing the influence of a common input signal. Our results demonstrate a causal link from the SC to the basal ganglia that could alter how sensory signals are used to guide perceptual choices without altering the sensory representations in visual cortex.

## Results

To measure the effects of superior colliculus (SC) inactivation on neuronal activity in the head of the caudate nucleus (CDh), we recorded the activity of CDh neurons while monkeys performed an attention task both before and during unilateral SC inactivation. In each experimental session, data were first collected over several blocks of trials before SC inactivation, muscimol was then injected into SC and the presence of a contralateral saccade deficit was confirmed, and finally data were collected over several additional blocks during the effects of inactivation (figure 1A). Because the SC output with access to caudate is almost totally ipsilateral (Grofov, 1979; Harting et al., 1980; Matsumoto et al., 2001; May et al., 2009; Nakano et al., 1990; Partlow et al., 1977), in a majority of sessions (n = 9) we inactivated SC on the “same-side” as recorded CDh neurons. We also collected data in several (n = 4) “opposite-side” SC inactivation plus recording sessions.

**Fig. 1.**
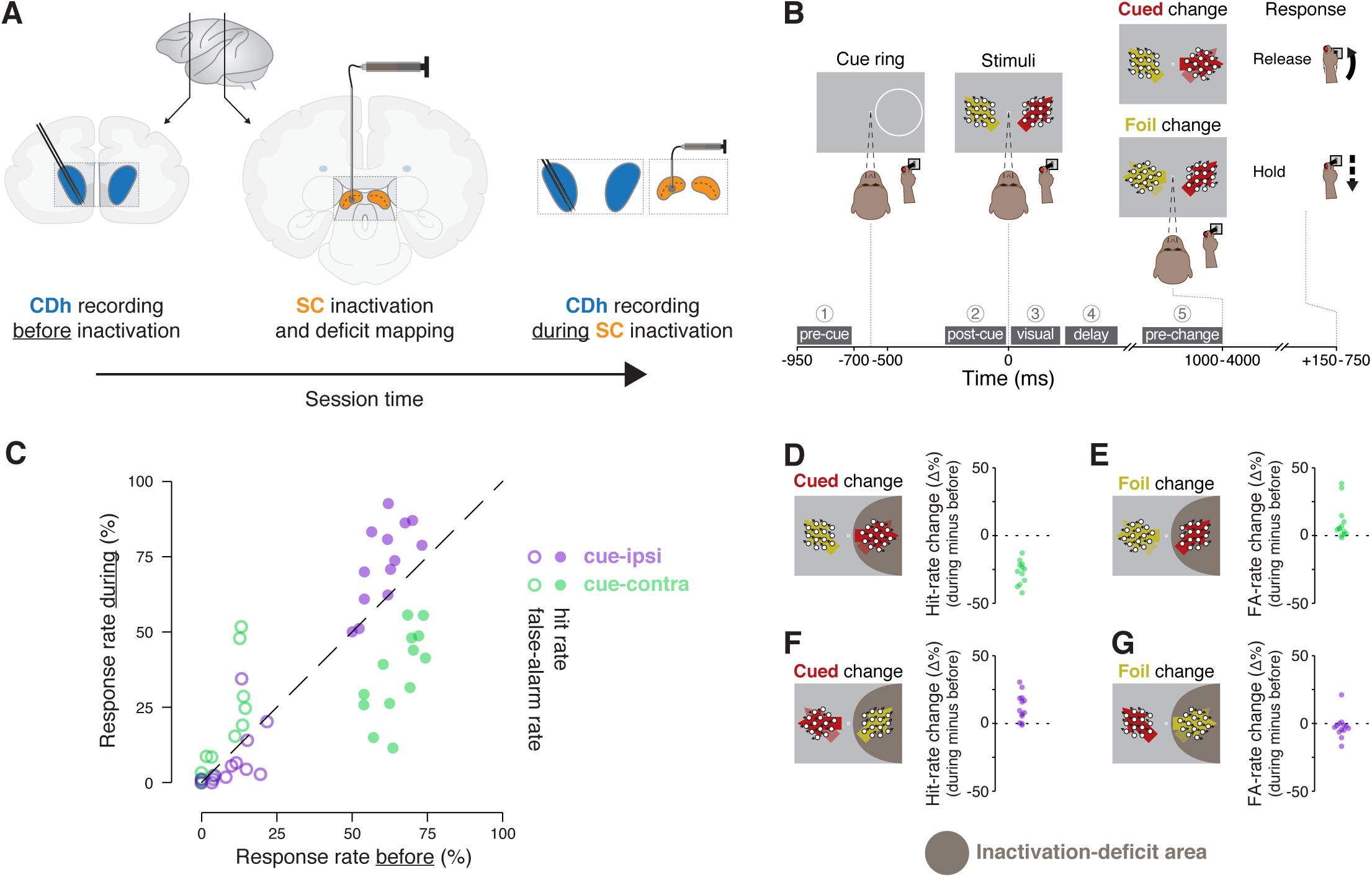
Experimental approach and behavioral effects of inactivation. **A**. In each session, the activity of neurons in the head of the caudate nucleus (CDh; blue shaded regions) was recorded with a pair of linear 32-channel probes before inactivation. An injection cannula was advanced into SC, 0.5ul of muscimol was infused, and the presence of a contralateral saccade deficit was confirmed (the images depict an inactivation of SC on the same-side of the brain as recorded CDh neurons; not depicted are opposite-side SC inactivations). A diagram of a lateral view of a macaque brain shows the antero-posterior positions (vertical black lines with adjoining arrows) corresponding to the diagrams of coronal slices displaying CDh and SC. During the effects of SC inactivation (referred to throughout as “during inactivation” or simply “during”), recordings of the activity of CDh neurons continued. **B.** Monkeys performed a cued motion-direction change detection task while CDh neuron activity was recorded before and during SC inactivation. Liquid reward was obtained obtained either by responding to a cued motion change with a joystick release (a hit), or by withholding response to a foil change (a correct reject); no reward was given if the monkey failed to respond to a cued change (a miss) or responded for a foil change (a false alarm). Dark-gray boxes with white text with circled numerals above show the timing, names, and numerical indexes of “task-epochs” used for data analysis. **C.** Performance summary comparing before and during SC inactivation. Hit (filled symbols) and false-alarm rates (empty symbols) during SC inactivation are plotted against rates before SC inactivation from the same sessions. Rates for cue-ipsilateral to SC inactivation site are plotted in purple and those for cue-contralateral are plotted in green. **D-G.** Differences in hit and false-alarm rates (during minus before SC inactivation) are plotted for each session, horizontal symbol spacing is artificially jittered to increase visibility. Brown shaded region depicts the area of visual space affected by muscimol inactivation of SC.

Neuronal and behavioral data were collected while monkeys performed a covert motion-change detection task. Two monkeys (P and R) were trained to release a joystick in response to a direction-of-motion change at a cued location and withhold their response if the change happened at a foil location (figure 1B). Monkeys obtained liquid reward either by responding to a cued motion change with a joystick release (a hit), or by withholding response to a foil change (a correct reject); no reward was given if the monkey failed to respond to a cued change (a miss) or responded to a foil change (a false alarm).

### Effects of SC inactivation on attention task performance

SC inactivation reliably produced spatially specifically impairments in attention task performance (figure 1C; S1). Consistent with previous reports (e.g. Zénon and Krauzlis, 2012), when the cue was presented inside the inactivation-deficit area of the visual field (cue-contra), hit rate decreased (figure 1D) and false-alarm rate increased (figure 1E) relative to performance before inactivation. When the cue ring was outside the deficit area (cue-ipsi), hit-rate increased (figure 1F) and false alarm-rate showed little change (figure 1G). We quantified the effects of inactivation on performance by comparing hit and false-alarm rates before to during SC inactivation with χ^2^-proportion tests (Fleiss et al., 2013), which confirmed that cue-contra hit rate during SC inactivation was significantly reduced in each session (13/13 sessions; all χ^2^ > 9.5, all p < 0.01). These tests also indicated that cue-ipsi hit rate increased significantly in 5/13 sessions (all χ^2^ > 7.1, all p < 0.01).

### Effects of SC inactivation on CDh spatial cue preferences

To examine the effects of SC inactivation on CDh neuronal activity, we used a pair of 32-channel linear probes in each inactivation plus recording session. This allowed us to make efficient use of each inactivation session, yielding 281 neurons identified as putative medium spiny neurons (MSNs, hereafter referred to as “CDh neurons”; 171 from monkey P and 110 from monkey R), collected across 13 inactivation sessions (9 same-side SC: 5 in monkey P and 4 in monkey R; 4 opposite-side SC: 2 each in monkeys P and R). MSN data from the two monkeys were pooled for subsequent analyses.

Many CDh neurons were robustly modulated by the location of the spatial cue, and these preferences changed during SC inactivation in different ways depending on whether we inactivated the same or opposite side. Before inactivation, an example neuron showed greater activity for cue presentation in the visual field contralateral to CDh recordings (cue-contra) over the ipsilateral visual field (cue-ipsi), illustrating a cue-contra preference (figure 2A). During inactivation of the same-side SC, a neuron recorded on the same contact (as in figure 2A) now exhibited a modest cue-ipsi preference (figure 2B). To quantify cue-side preferences of individual neurons, we computed the area under a receiver operating characteristic curve (ROC area; Britten et al., 1992) in each of several temporal epochs defined by task events (1: pre-cue, 2: post-cue, 3: visual, 4: delay, 5: pre-change; figure 1B). Before same-side SC inactivation, the example unit (in 2A) had significant cue-contra preferences in epochs 1-4 (figure 2C; all bootstrapped 95% confidence intervals greater than 0.5). During same-side SC inactivation, the example unit (in 2B) had no significant cue-side preference in epochs 1-4 (all 95% ROC CIs ⊄ 0.5), and a significant cue-ipsi preference in epoch 5 (95% ROC CI < 0.5; figure 2D). Thus same-side SC inactivation appeared to markedly reduce the cue-contra preference.

**Fig. 2.**
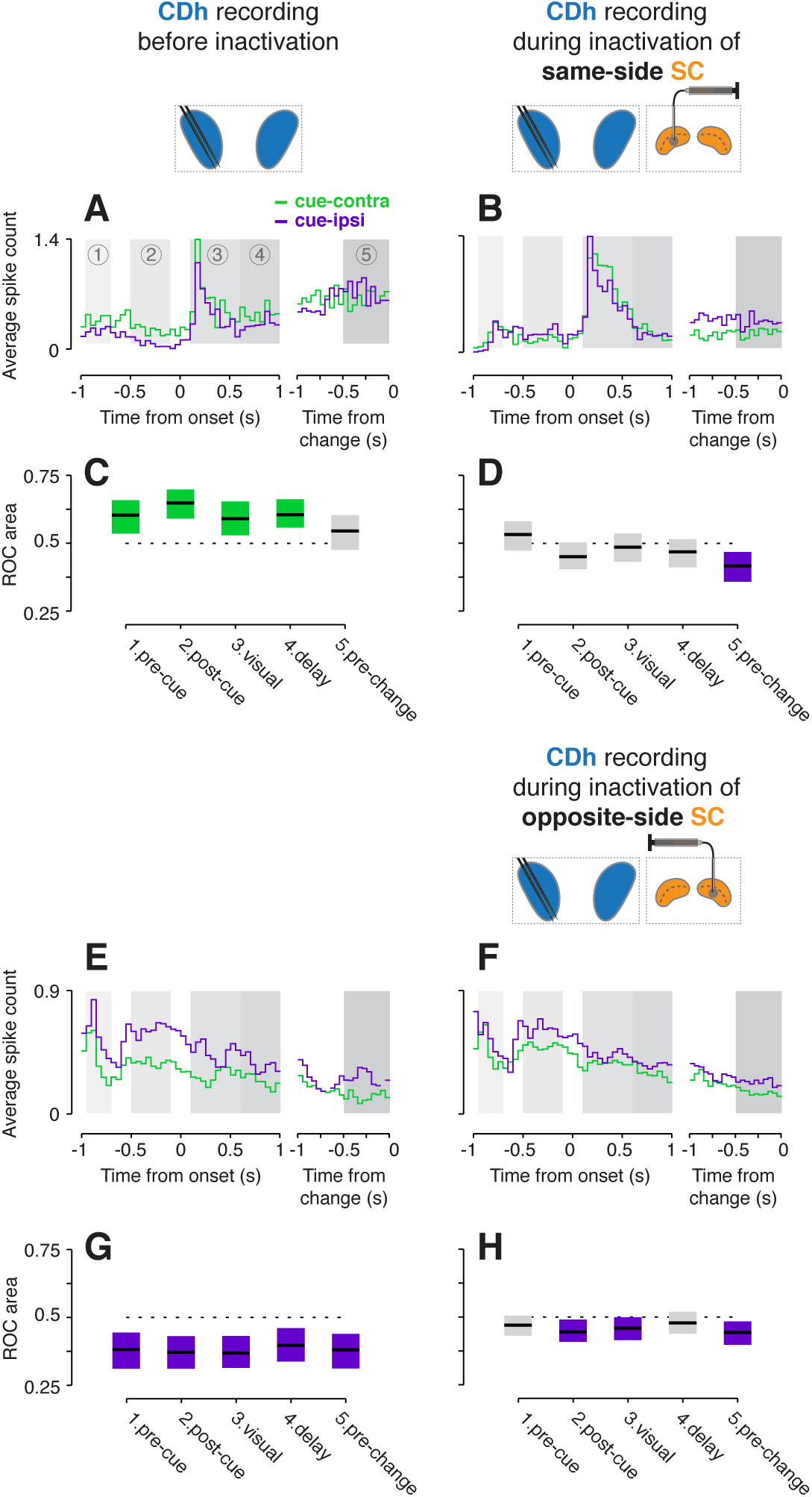
Example CDh neuron activity before and during SC inactivation. **A.** Activity of an example CDh neuron recorded before same-side SC inactivation. Traces representing spike counts in non-overlapping 50ms bins are plotted separately for cue-contralateral (to CDh recordings) and cue-ipsilateral conditions. A portion of data aligned on stimulus onset is presented in the left panel, and a portion aligned on stimulus change in the right panel. Shaded areas mark five task-epochs: 1: pre-cue, 2: post-cue, 3: visual, 4: delay, 5: pre-change. **B.** Presentation as in panel **A** but for an example unit recorded *during* same-side SC inactivation. This example unit was recorded on the same electrode contact as the unit in panel **A. C.** The area under a receiver operating characteristic curve (ROC area) was used to quantify the cue-side preference of the example unit in **A** in each task-epoch. Horizontal black line segments are ROC areas and surrounding shaded regions indicate bootstrapped 95% confidence intervals (CIs). Individual ROC areas are considered significant when their 95% CIs ⊄ 0.5 (do not contain 0.5; dotted line); significant cue-contra ROC CIs are colored green and significant cue-ipsi CIs are colored purple, gray ROC CIs indicate non-significant cue-side preferences. **D.** ROC areas and 95% CIs comparing cue-contra to cue-ipsi for the example neuron in panel **B**. Presentation as in panel **A** but for an example unit recorded before opposite-side SC inactivation. **F.** Presentation as in panel **E** but for *during* inactivation. This example unit was recorded on the same electrode contact as the unit in panel **E. G.** ROC areas and 95% CIs comparing cue-contra to cue-ipsi for the example neuron in panel **E. H**. ROC areas and 95% CIs comparing cue-contra to cue-ipsi for the example neuron in panel **F**.

In contrast, inactivation of the opposite-side SC produced small decreases in the number of CDh neurons with ipsi-cue preference. Another example neuron recorded in a separate session had significant cue-ipsi preferences in all epochs (figure 2E; all ROC CIs < 0.5). During opposite-side SC inactivation, a neuron on the same contact (as in figure 2E) showed weaker cue-ipsi preferences (figure 2F) which remained significant in epochs 2, 3 and 5 (figure 2H, all ROC Cis < 0.5). Together, these examples illustrate the overall pattern in our results – unilateral SC inactivation reduced cue-side preferences, and these reductions were largest for CDh neurons with contra-cue preferences recorded on the same side as the inactivation.

Across our population of CDh neurons, same-side SC inactivation redistributed the cue-side preferences in favor of cue-ipsi and opposite-side SC inactivation weakly pushed preferences towards cue-contra. Before same-side SC inactivation (figure 3A), 24.5% of CDh neurons had significant cue-contra preferences and 15.1% had significant cue-ipsi preferences (figure 3E; all ROC CIs ⊄ 0.5). During same-side SC inactivation (figure 3B), just 9.7% had significant cue-contra preferences and 24.7% had significant cue-ipsi preferences (figure 3F; all ROC CIs ⊄ 0.5; all percentages collapsed across epochs). In contrast, opposite-side SC inactivation increased the prevalence of significant cue-contra preferences from 12.7% to 17.5% and decreased cue-ipsi from 16.7% to 12.5% (figure 3C, D, G, H; all ROC CIs ⊄ 0.5).

**Fig. 3.**
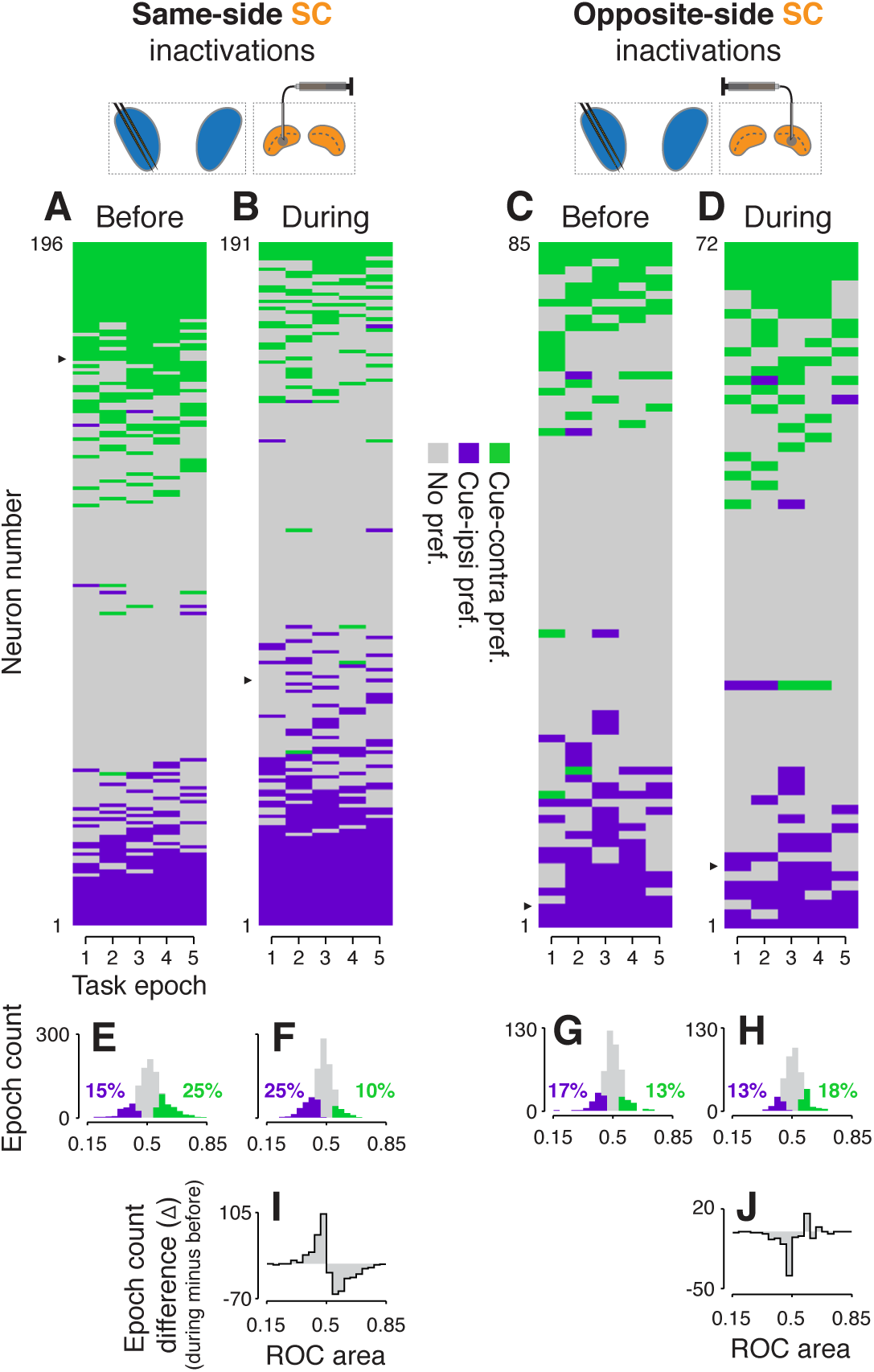
Effects of SC inactivation on CDh population cue-side preferences. **A**. Cue-side preferences in each task-epoch (columns) for each CDh neuron (rows) recorded before same-side SC inactivation. Green cells indicate significant cue-contra preference, gray cells indicate no significant preference, and purple cells indicate significant cue-ipsi preference. Rows (neurons) are sorted from most cue-contra preference at top to most cue-ipsi preference at bottom. Black arrowhead at left edge indicates the example neuron shown in figure 2A. **B**. Presentation as in panel **A**, but for neurons recorded *during* same-side SC inactivation. Black arrowhead at left edge indicates the example neuron shown in figure 2B. **C**. Presentation as in panel **A**, but for neurons recorded before opposite-side SC inactivation. Black arrowhead at left edge indicates the example neuron shown in figure 2E. **D**. Presentation as in panel **A**, but for neurons recorded *during* opposite-side SC inactivation. Black arrowhead at left edge indicates the example neuron shown in figure 2F. **E**. Histogram of ROC areas comparing cue-contra to cue-ipsi for all CDh neurons recorded before same-side SC inactivation, collapsed across task-epochs. Green bars indicate significant cue-contra preferences, and purple bars indicate significant cue-ipsi preferences. Colored text indicates percentages of significant cue-side preference epochs. **F**. Presentation as in **E**, but for during same-side SC inactivation. **G**. Presentation as in **E** but for before opposite-side SC inactivation. **H**. Presentation as in **E** but for *during* opposite-side SC inactivation. **I**. Difference-of-histograms plot showing the change in ROC area distribution during same-side SC inactivation. Histogram values in **E** were subtracted from values in **F**, ignoring cue-side preferences. **J**. Presentation as in **I** but for opposite-side SC inactivation data.

To statistically test the effects of SC inactivation on the prevalence of single-neuron cue-side preferences, we fit proportions of significant cue-contra and cue-ipsi preferences in each task epoch, before and during inactivation, with logistic regression (separately for same-side and opposite-side SC inactivation data). This analysis confirmed that same-side inactivation significantly reduced the proportion of cue-contra preferring CDh units (tStat = −3.2336, p < 0.01) and increased the proportion preferring cue-ipsi (interaction term; tStat = 4.2445, p << 0.01) whereas opposite-side inactivation had no significant effect on the proportion of units preferring either cue-contra (tStat = 0.5689, p = 0.57) or cue-ipsi (interaction term; tStat = −1.1691, p = 0.24). In addition, both regression models indicated no effect of task epoch (same-side model tStat = 0.3151, p = 0.75; opposite-side model tStat = −1.3342, p = 0.18).

A difference in ROC area distributions (during minus before; figure 3I) illustrates that during same-side SC inactivation, the reduction of significant cue-contra preferring neurons was accompanied by an increase in non-significant cue-ipsi preferences; the reverse trend is evident for opposite-side SC inactivation (figure 3J). Quantitatively, same side SC inactivation caused the skewness of the ROC area distribution to change, significantly, from positive to negative (before: skewness = 0.12, 95% bootstrapped CI = [-0.05, 0.29]; during: skewness = −0.29, 95% CI = [-0.47, −0.11]; p < 0.05; bootstrap test), and caused a small but significant change in the distribution’s mean (before: μ = 0.515, 95% bootstrapped CI = [0.509, 0.520]; during: μ = 0.478, 95% CI = [0.474, 0.482]; p < 0.05, bootstrap test). Thus, the categorical shift in cue-side preferences caused by same-side SC inactivation was due to a redistribution of preferences, not just an overall translation of the distribution of ROC areas.

In summary, these findings demonstrate that the modulation of CDh neuronal activity by spatial cues depends on activity from the SC on the same side of the brain.

### Changes in spike-count correlations of CDh neurons

We next examined pairwise spike-count correlations to assess whether SC inactivation altered the influence of some common input signal to CDh (Cumming and Nienborg, 2016). We computed correlations in all combinations of task epoch, cue-side condition, and inactivation-state (before / during). Values were pooled across epochs and cue-side because there were no significant differences across these conditions (bootstrap tests, smallest p = 0.21). We then fit the resulting distributions simultaneously (see Methods) and used the fits to determine whether inactivation had changed the shape of the distributions (figure 4). Fitting showed that the distribution of correlations during same-side SC inactivation (figure 4C) was significantly narrower than before (figure 4A; bootstrap test, p << 0.01); the width parameter of the fitted Stable distribution narrowed from 0.071 (95% bootstrapped CI = [0.07, 0.072]) to 0.057 (95% CI = [0.056 0.058]). A difference of density histograms (during minus before) illustrates that same-side inactivation caused a reduction in both positively and negatively correlated pairs, and an increase in weakly correlated and uncorrelated pairs (figure 4E). In contrast, opposite-side SC inactivation caused no change in the width of correlation distributions (figure 4B, 4D; bootstrap test, p = 0.88; fitted Stable distribution width parameter before = 0.062, 95% CI = [0.061, 0.063]; during = 0.061, 95% CI = [0.06, 0.062]), but did result in a significant increase in skewness from 0.76 (95% bootstrapped CI = [0.64, 0.97]) to 1.88 (95% CI = [1.68, 2.09]), which can be seen in the small increase of positive values in the during minus before difference of density histograms (figure 4F). Importantly, because pairs of neurons with lower firing rates tend to be less strongly correlated (Cohen and Maunsell, 2009; Ecker et al., 2010; la Rocha et al., 2007; Mitchell et al., 2009), we compared firing rates before SC inactivation (same-side before: μ = 9.66 spikes/s, bootstrapped 95% CI = [9.57, 9.75]; opposite-side before: μ = 9.39 sp/s, 95% CI = [9.27, 9.52]) to rates during (same-side during: μ = 9.57 sp/s, 95% CI = [9.47, 9.66]; opposite-side during: μ = 9.42 sp/s, 95% CI = [9.27, 9.52]) and found no significant effect of inactivation on firing rate within sessions (smallest tStat = −0.6715, smallest p = 0.5) or across sessions (same-side: tStat = 0.6764, p = 0.49; opposite-side inactivations: tStat = 2.1392e-12, p = 1).

**Fig. 4.**
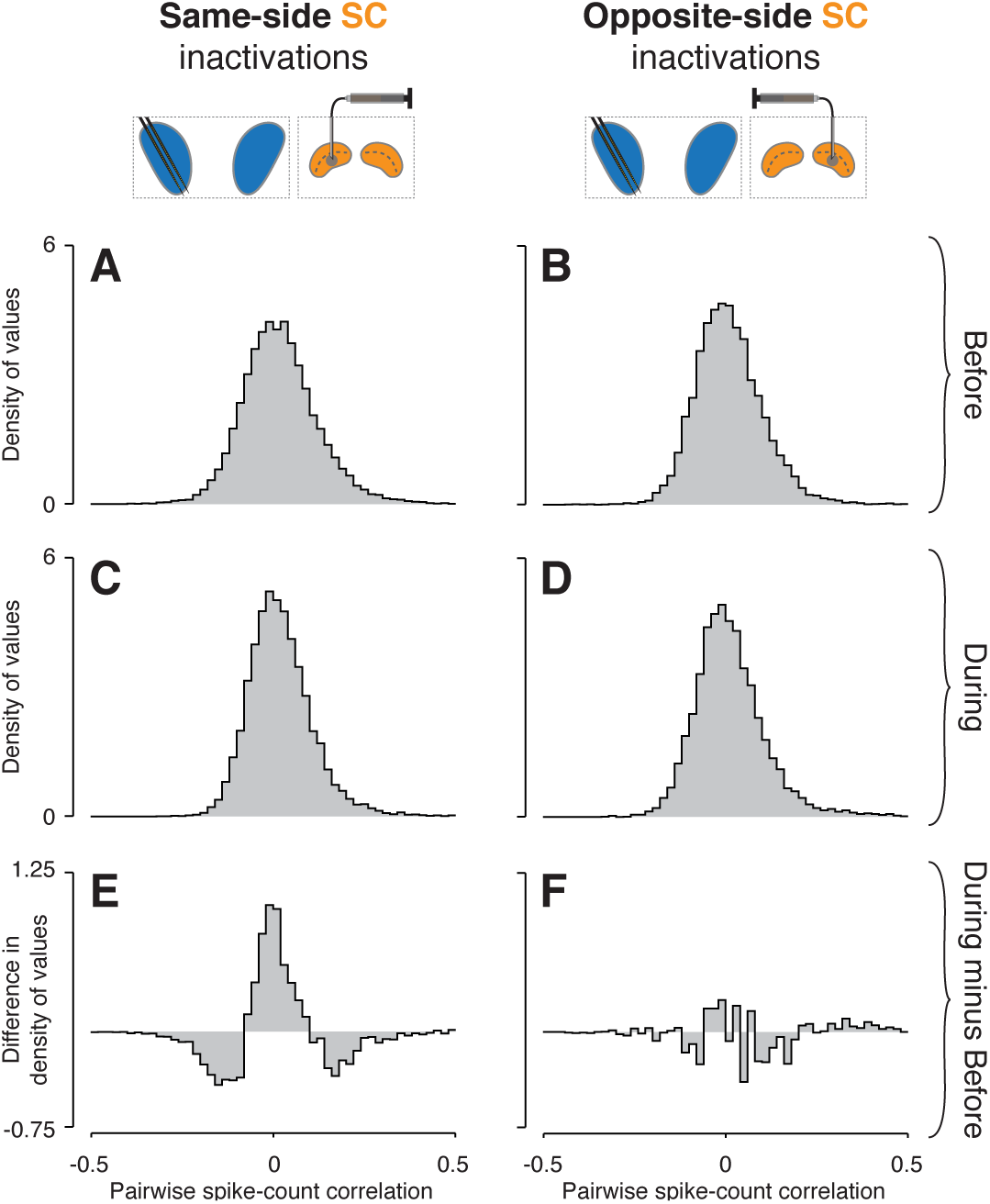
Inactivation effects on CDh pairwise neuronal correlations. **A.** Before same-side SC inactivation density histogram of pairwise spike-count correlations pooled across epochs and cue-side presentations (n = 4280 pairs). **B.** Presentation as in **A** but for before opposite-side SC inactivation (n = 2302 pairs). **C.** Presentation as in **A** but for *during* same-side SC inactivation (n = 4042 pairs). **D**. Presentation as in **B** but for *during* opposite-side SC inactivation (n = 1468 pairs). **E**. Density histogram difference: during same-side SC inactivation minus before same-side SC inactivation. **F.** Presentation as in **E** but for opposite-side SC inactivations.

From these results, we conclude that same-side SC inactivation reduces the influence of some common input to CDh. Because removing this common influence results in both fewer positively and fewer negatively correlated pairs, it must ultimately have excitatory effects on some CDh neurons and inhibitory effects on others.

### Changes in classifier performance based on CDh neuronal activity

To test how SC inactivation altered the information content in our recorded CDh populations, we examined the performance of multi-dimensional classifiers, which rely on the information contained in the population-level structure of neuronal activity. Specifically, we asked how well CDh population activity encoded information about the cue-side and task-epoch, and whether SC inactivation affected that encoding.

For each session, we trained and tested two classifiers: the first with CDh neuronal activity from before SC inactivation and the second with activity recorded during inactivation. Each classifier took an n-dimensional vector of neuronal activity (where n is the number of simultaneously recorded neurons before or during inactivation) from one epoch in one trial and returned a “classifier epoch index” that uniquely identified the task epoch and cue-side (figure 5A). Classifier epoch index duplicated task-epochs 1-5 to include 1-5 cue-contra and 1-5 cue-ipsi, and was compared to the “true epoch index” (the task-epoch and cue-side from which the data actually arose) to generate a confusion matrix (figure 5B) showing how frequently each classifier epoch index was correctly identified. To help interpret the confusion matrix, we also highlighted three categories of classification outcome relevant to our specific scientific questions: (1) correct classifications, (2) cue-side misclassifications, and (3) epoch misclassifications (figure 5B). After comparing the performance of several classifier variants, we found that a “boosted decision tree” yielded the best correct-classification rates over all datasets and used this variant for subsequent analyses.

Same-side SC inactivation impaired the ability of classifiers to decode both cue-side and task-epoch information. Before same-side SC inactivation, correct classification rates were high for all epochs on both cue-sides (figure 5C), in aggregate 70% across sessions and epoch indexes (figure 5F). High aggregate classifier performance resulted from largely consistent patterns of performance in individual same-side SC inactivation sessions (figure 5F). During same-side SC inactivation, classification performance was poor across epoch indexes (figure 5D), with correct-classification rates falling to about 47% (figure 5G).

**Fig. 5.**
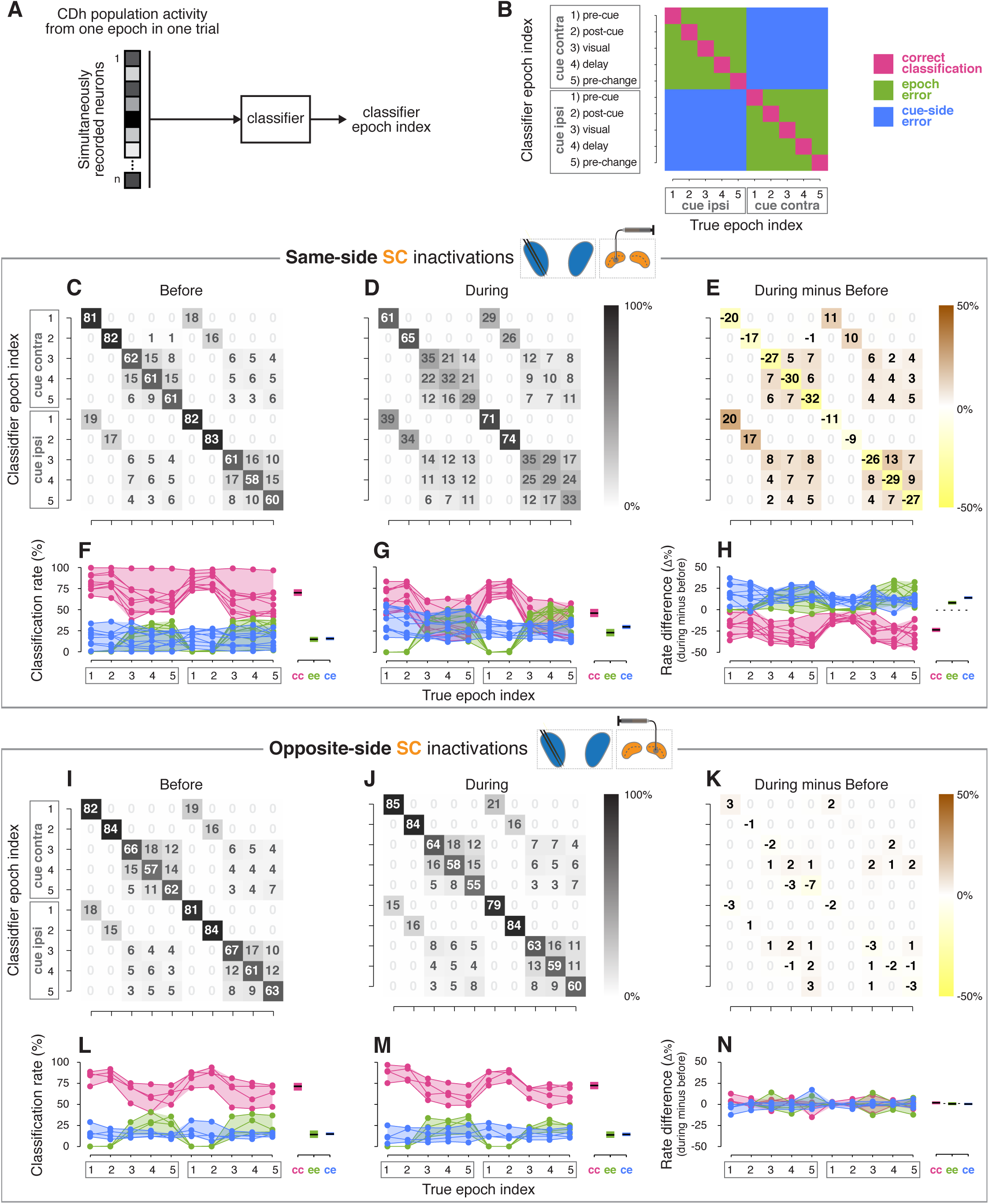
Classifier analyses. **A.** A boosted decision tree classifier took n-dimensional vectors (where n is the number of simultaneously recorded neurons before or during SC inactivation) of CDh neuronal activity from individual epochs in single trials and returned a “classifier epoch index” indicating the classifier’s guess regarding both which task-epoch and which cue-side condition the activity came from. **B.** A confusion matrix is traditionally used to show how frequently each possible classifier epoch index (vertical axis) is applied for each true epoch index. Here we have additionally color-coded a confusion matrix to highlight three categories of classification outcome: (1) correct classifications (magenta), (2) cue-side misclassifications (blue), and (3) epoch misclassifications (green). **C.** Across-sessions aggregated cross-validated confusion matrix before same-side SC inactivation. Cell shading starts at white for 0% with larger values grading darker up to black at 100%. For each “true epoch index” column (for example, “pre-cue cue-contra” in column 1), the numeral in each row is the (rounded) percentage of times classifiers assigned that row’s class label to all inputs of that column’s input (true) class. For example, the number 81 in the 1st row and 1st column indicates that, across classifiers / sessions, when classifiers were given “pre-cue cue-contra” input vectors, 81% of those vectors were labelled as “pre-cue cue contra”; the number 19 in column 1, row 6 indicates that the remaining 19% of the time, classifiers erroneously labelled “pre-cue cue contra” activity vectors as “pre-cue cue-ipsi”. **D**. Presentation as in **C** but for *during* same-side SC inactivation. **E**. Difference of aggregated confusion matrixes (during minus before). Cell color mapping starts at saturated yellow for −50% with larger values grading to white at 0%, followed by increasing positive values grading up to saturated brown at +50%. **F.** Left: per-session breakdown of classifier performance with classification rates divided into outcome categories shown in panel **B**. Connected dots are from a single session. For each true epoch index on the horizontal axis, colored dots show the percentage of times the single-session classifier epoch index fell into one of the three classification categories defined in **B** (correct classification, epoch misclassification, cue-side misclassification). Right: per-category breakdown of mean classification rates across sessions and true epoch indexes; cc: correct classifications, ee: epoch errors, ce: cue-side errors. Shaded areas indicate 95% bootstrapped confidence intervals on mean from individual session classification rates. **G.** Presentation as in panel **F**, but for *during* same-side SC inactivations. **H**. Left: per-session differences in classification categories (during minus before). Right: per-category breakdown of difference in mean classification rates across sessions and true epoch indexes. Dotted line marks 0. **I-N**. Presentation as in panels **C-H** but for opposite-side SC inactivation sessions.

To highlight the effects of SC inactivation on classifier performance, we examined a difference of confusion matrices (during minus before; figure 5E). Cue-contra epoch indexes were slightly more strongly affected than cue-ipsi, with correct classification rates falling by 26% for cue-contra and 21% for cue-ipsi (figure 5H). Misclassification errors induced by same-side SC inactivation were more frequently cue-side errors, which rose by 16%, than epoch errors, which rose by 9% (figure 5H). To statistically test changes in classification rates caused by inactivation, we used logistic regression on rates with session, classification-outcome category, and true epoch index as predictors. This analysis confirmed that correct-classification rates were significantly poorer during same-side SC inactivation compared to before (tStat = −2.3893, p < 0.02), and that cue-side misclassification rates were more pronounced than epoch misclassification rates (interaction term: tStat = 2.2681, p < 0.03).

Opposite-side SC inactivation had essentially no effect on classification performance. Before opposite-side SC inactivation (figure 5I), the aggregate correct-classification rate was 71% (figure 5L) and rose to 72% during inactivation (figure 5J, M). Again, these patterns of performance were consistent across sessions (figure 5L-M). Logistic regression applied to data from these opposite-side SC inactivation sessions revealed no significant effect of inactivation (tStat = −0.54726, p = 0.5842), and no significant interactions (tStat range: −0.05 - 0.075, all p > 0.91).

From these classifier-based analyses we conclude that same-side SC inactivation primarily disrupts CDh population encoding of the task-relevant spatial location, and also interferes with non-spatial encoding of task epochs.

## Discussion

Our results establish that the pattern of neuronal activity observed during attention-task performance in the anterior portion or “head” of the caudate nucleus (CDh) relies on output from the superior colliculus (SC), and provide a possible basis for the reliable, performance-altering effects of SC inactivation during attention tasks. Inactivation of SC on the “same-side” of the brain as recorded CDh neurons clearly and consistently disrupted attention-related modulation in CDh whereas inactivation of “opposite-side” SC did not. Specifically, SC inactivation redistributed the cue-side preferences of individual CDh neurons, decreasing the prevalence of cue-contra preferences and increasing cue-ipsi preferences. Inactivation also disrupted a non-spatial component of CDh activity, impairing the ability of a classifier to correctly decode task-epoch from populations of CDh neurons. Inactivation of the same-side SC also narrowed the distribution of spike-count correlations between pairs of CDh neurons, consistent with the interpretation that the SC normally provides a shared input to CDh neurons that was reduced during inactivation. These results provide causal evidence that a circuit mechanism from the SC to the basal ganglia is part of the control system for covert visual attention.

### A reversal of the classic subcortical hierarchy

Our demonstration that SC output strongly influences CDh is a reversal of the typically depicted subcortical hierarchy in which information flows from cortex through the basal ganglia to the SC (e.g. Hikosaka et al., 2014). This adds a recurrent or ascending component to the already well-established circuit by which CDh output influences SC (Yasuda and Hikosaka, 2015). According to the classic circuit diagram, caudate output, acting through the external segment of the globus pallidus (GPe) and substantia nigra pars reticulata (SNr), modulates the excitability of SC neurons by removing or adding inhibition, thereby affecting saccade probability (Hikosaka et al., 2006). This mechanism can help primates orient towards visual stimuli associated with large rewards and away from those associated with small rewards (Amita and Hikosaka, 2019; Yamamoto et al., 2013).

Our results complement this picture by demonstrating the importance of signals arising from the SC and sent back to the basal ganglia. There are two routes by which neurons in the intermediate and deep layers of SC – which our inactivations targeted – might affect CDh neuronal activity: (1) through the substantia nigra pars compacta (SNc), or (2) through the centre médian-parafascicular complex (CM-Pf) of the thalamus (Krauzlis et al., 2013). Previous experimental work in monkeys has indicated both of these pathways from SC to CDh might be involved in the control of spatial attention.

With regard to the first route, dopaminergic neurons in SNc can be activated by SC and broadcast signals widely throughout the striatum that are necessary for normal processing of visual information. In monkeys with V1 lesions, responses of SNc DA neurons to reward-predicting visual stimuli are virtually eliminated by SC inactivation (Takakuwa et al., 2017), demonstrating that primate SC can excite SNc DA neurons. When SNc DA signals to caudate are cut off by unilaterally infusing MPTP into caudate, the result is contralateral visual hemineglect (Miyashita et al., 1995). Monkeys in this state are still able to make contraversive saccades when presented with single targets but fail to do so under free viewing conditions or when presented with a pair of lateralized saccade choice targets. These results are consistent with the contralateral attention deficits caused by SC inactivation being partly mediated by a circuit through SNc to caudate.

The second route, from SC through the CM-Pf complex to CDh, has been implicated in the control of attention in both monkeys and humans. Neurons in the intermediate and deep layers of SC send a predominantly ipsilateral projection to the CM-Pf (Harting et al., 1980; Partlow et al., 1977) which, in turn, sends an exclusively ipsilateral projection to the striatum (Matsumoto et al., 2001; Nakano et al., 1990). Unilateral muscimol inactivation of CM-Pf in monkeys removed the reaction time (RT) benefit conferred by a spatial cue in an attention task, but only for the contralateral hemifield not the ipsilateral field (Minamimoto and Kimura, 2002). In humans performing a spatial attention task, fMRI data indicated that the CM-Pf was consistently activated by the “attentional shifts” required in that task (Hulme et al., 2010). These data are consistent with the SC transmitting signals related to attentional selection of contralateral visual stimuli to caudate through CM-Pf.

Whether via one of these circuits or some other pathway, our finding that same-side SC inactivation strongly alters CDh neuron cue-side preferences establishes the importance of recurrent interactions between the SC and basal ganglia during covert visual attention tasks. This interaction includes modulation of neuronal excitability in SC by basal ganglia output and shaping of spatial selectivity in caudate neurons by SC. Our proposal for the functional role of this “subcortical loop” differs from previous conceptions (McHaffie et al., 2005; Redgrave et al., 2010; Redgrave and Gurney, 2006) in the primary emphasis we place on a visual selection mechanism. An abundance of evidence has accumulated for the SC’s causal participation in visual selection – SC activity helps determine which visual information is used to guide perceptual reports (Cavanaugh and Wurtz, 2004; Herman et al., 2018; Lovejoy and Krauzlis, 2010; Müller et al., 2005; Zénon and Krauzlis, 2012). Attention is impaired when SC is inactivated, or when the nodes allowing SC activity to reach caudate are lesioned. Together with the CDh neuronal correlates of SC inactivation we observed, we consider this strong evidence that disruption of the recurrent interactions between SC and basal ganglia are a major reason why SC inactivation causes deficits in covert visual attention.

### Task-state representation

Our classifier results are compatible with striatal circuits representing “belief states” that summarize environmental conditions. Many previous studies have found that striatal neurons are active at distinct times during a task. The activation of unique subsets of striatal neurons at distinct times has been variously interpreted to mean this activity represents action sequences (Jog et al., 1999; Kermadi and Joseph, 1995; Miyachi et al., 2002; Sheng et al., 2019), accumulated evidence (Ding and Gold, 2010), uncertainty about object-reward associations (White and Monosov, 2016), stimulus or reward expectation (Hikosaka et al., 1989), or action value (Lau and Glimcher, 2008; Lee et al., 2015; Samejima et al., 2005; Seo et al., 2012). We consider the heterogenous joint representation of multiple task- and internal-state variables displayed by striatal neurons across experimental conditions as most consistent with striatal population encoding of “belief state” (Daw et al., 2006; Rao, 2010). Belief states provide an elegant way to apply reinforcement learning (RL) methods to the problem of learning environmental state-contingent action values in the face of uncertainty about environmental state, which aligns with inherently stochastic sensory representations found in the brain (Gershman and Uchida, 2019).

Striatal encoding of belief state is compatible with our classifier results and offers a novel interpretation of changes in attention-task behavior resulting from SC inactivation. If striatal neurons represent belief state, it should be possible to accurately decode environmental conditions from the activity of those neurons, as our classifier results demonstrated. The defining features of our attention-task space (cue-side and task-epoch) amount to the task’s “environmental conditions”, which we found can be decoded from CDh neuronal activity with high accuracy (∼70%) before SC inactivation (figure 5; Arcizet and Krauzlis, 2018). The impaired decoding of task-state observed during same-side SC inactivation then implies a disorganized underlying belief state representation. If SC inactivation degrades belief state representation, the behavioral effects of SC inactivation during attention tasks amount to errors in estimating the value of responding or withholding responses for task events. In contrast to standard views of attention, which focus on the quality of sensory representations, this interpretation suggests that some of the behavioral and neurophysiological correlates of attention can be understood as consequences of mechanisms for learning the value of actions or the value of state-contingent action values (sensory-motor associations). From this perspective, SC inactivation reduces hit-rate by reducing the subjective value of responding to cued changes and increases false alarm-rate by inflating the subjective value of responding to foil changes.

Our assertion that some correlates of attention arise from learning mechanisms is bolstered by the striking correspondence between sub-cortical structures necessary for attention and those implicated in learning. In monkeys that have learned to associate a “conditioned stimulus” (CS) with either an appetitive or an aversive stimulus, tonically active neurons (TANs) in the striatum exhibit stereotypical “pauses” in spiking for visual or auditory CS presentation (Aosaki et al., 1994b), but not for unpaired stimuli (for which no association has been learned). These learning-induced responses of striatal TANs require both intact SNc DA projections to striatum (Aosaki et al., 1994a), and normal signaling from CM-Pf (Matsumoto et al., 2001). Striatal TANs are thought to be large aspiny cholinergic interneurons which can bidirectionally regulate the influence of cortical input to MSNs pre- and post-synaptically (Ding et al., 2010). These results show that visual stimulus-related signals flowing through SNc and CM-Pf to the striatum have the potential to alter the sensitivity of inputs to striatal MSNs by affecting the activity of striatal TANs. The same routes through SNc and CM-Pf that likely allow SC to influence attention-related activity in caudate are also necessary for expression of learning-related activity changes in striatum.

## Conclusions

We have provided causal evidence that attention-related modulation in caudate neurons depends on output from the superior colliculus. Ascending signals from the SC through one or more possible circuits are also necessary for CDh population representation of task-state variables. These results offer a novel basis for attention-task deficits during SC inactivation as the result of altered processing of stimulus relevance in the basal ganglia and demonstrate the general importance of circuits through the basal ganglia for the control of spatial attention.

## Methods

### Animals

Data were collected from two adult male rhesus macaques (P and R; *Macaca Mulatta*) weighing 10-16 kg. All experimental protocols were approved by the National Eye Institute Animal Care and Use Committee, and all procedures were performed in accordance with the United States Public Health Service policy on the humane care and use of laboratory animals.

### Tasks & Stimuli

The details of our apparatus, task, and stimuli have been described in detail previously (Arcizet and Krauzlis, 2018), but are partially reproduced here for clarity. Monkeys were seated in primate chairs (Crist Instrument, Hagerstown, MD, United States) with head fixed, inside a darkened booth. Animals were positioned with eyes 48 cm from an LCD display with a refresh-rate of 100 Hz (VIEWPixx; VPixx Technologies, Saint-Bruno, QC, Canada), and experiments were orchestrated with a modified version of PLDAPS (Eastman and Huk, 2012), running on a “MacPro5,1” (Apple Inc., Cupertino, CA). Eye position was monitored using an EyeLink 1000 infrared eye-tracking system (SR Research, Ottawa, Ontario, Canada), and manual responses were collected with a single-axis joystick (CH Products, model HFX-10) mounted to the primate chair and oriented to allow vertical movement.

Monkeys initiated each trial by pressing down on a joystick which triggered the appearance of a central fixation square. Central gaze fixation within a 2-3° window was required for the entire duration of the trial; exiting the window caused the trial to be aborted and repeated. After an initial 250 ms of maintained fixation, a cue-ring (inner radius 3.75°, outer radius 4°) was presented for 200 ms at an eccentricity of 10° (figure 1B). Two motion-dot patches (3° radius) were presented 500ms after the cue-ring was extinguished; the “cued” patch was centered on the former location of the cue-ring, and the “foil” patch was presented at an equally eccentric opposing location (180° of elevation away). The location of the spatial cue was constant for a block of 68 trials and then switched to the opposite location. A motion-direction change was possible 1000 ms – 4000 ms following stimulus onset (uniform distribution), and stimuli were extinguished 1000 ms after the change (i.e., maximum stimulus duration was 5000 ms). If the cued stimulus changed, the monkey was required to release the joystick in a time window 200 ms - 800 ms after the change; if the foil stimulus changed the monkey was required to maintain joystick press for an additional 1000ms. If the monkey released the joystick for a cued change (“hit”) or maintained joystick press for a foil change (“correct reject”), a liquid reward was delivered 1000 ms after the change; no reward was delivered for failing to release the joystick for a cued change (“miss”) or releasing the joystick for a foil change (“false alarm”). In each trial, either the cued stimulus changed, or the foil stimulus changed, but not both.

Because of the idiosyncratic and heterogeneous quality of the stimulus-placement preferences of caudate neurons we observed previously (Arcizet and Krauzlis, 2018), we chose stimulus locations to ensure behavioral effects of superior colliculus (SC) inactivation. The two diametrically opposed stimulus locations were chosen at the start of each session on the basis of the planned inactivation-deficit area so that, during SC inactivation, one of the two stimuli would fall inside the deficit area and one outside.

The direction of motion of dots in each stimulus patch was drawn from gaussian distributions with standard deviation s = 16°. The magnitude of the motion-direction change was adjusted from session-to-session to maintain relatively constant performance and was generally kept in a range of 13° - 22°. The mean of the distribution for the cued stimulus patch and the foil stimulus patch varied from day to day but always differed by 90°. Each dot was 6 pixels in diameter, moved at a speed of 15 degrees per second, had a lifetime of 100 ms (10 frames), and overall dot density was 26 per square degree per second.

The delayed visually guided saccade task used to assess the effects of muscimol injection into SC has been described previously (Zénon and Krauzlis, 2012).

### Neuronal Recordings & Inactivation

At the start of each session, two 32-contact “v-probes” (200 μm spacing between contacts in a single-column geometry; Plexon Inc., Dallas, TX) were advanced into the left caudate nucleus “head” (CDh), and one injection canula was advanced into cortex overlying SC; both probes and injection canula were controlled with a micromanipulator (NAN Instruments LTD, Nof Hagalil, Israel). CDh recording sites ranged from AC+4 to AC+8, where AC refers to the anterior-posterior location of the anterior commissure, located at approximately AP20 relative to ear-bar zero with structural MRI images. We localized recording contacts to the CDh based on position information from structural MRI and on the basis of the low background activity observed on the contacts.

CDh neuronal data were collected over 6-8 blocks (408-544 trials) of attention task performance before SC inactivation. Once before-inactivation data collection was complete, the injection canula was advanced to an estimated depth of 2.5 mm below the dorsal surface of the SC, at which depth 0.5 μL of 5 μg/μL muscimol (a GABA_A_ agonist) was injected. Following muscimol injection, peak velocities of visually guided saccades to a variety of locations in the visual field were measured and compared to those from previous (non-inactivation) sessions to confirm the presence and spatial extent of a deficit in the contralateral visual field. In 9/13 “same-side” SC inactivation sessions (5 in monkey P and 4 in monkey R), muscimol was injected into left SC, and in the remaining 4/13 “opposite-side” inactivations (2 in monkey P and 2 in monkey R), the injection was into right SC. Once a saccade deficit was confirmed, during-inactivation CDh neuronal data were collected over 8-14 blocks of the attention task. Following collection of CDh neuronal data in the attention task during-inactivation, visually guided saccades were again used to map the spatial extent of the inactivation-deficit area.

### Spike Sorting & Unit Selection

Raw voltage signals from each v-probe contact was digitized (40kHz sample-rate), high-pass filtered and stored with an “Omniplex D” system (Plexon Inc., Dallas, TX). These “continuous spike channel” data were analyzed offline with Kilosort (Pachitariu et al., 2016), including manual curation, to identify putative single neuron waveforms. Our intention was to identify and analyze spike data from caudate “medium spiny projection neurons” (MSNs) also called “phasically active neurons” (PANs). To identify MSNs, we followed previously established methods based on each putative neuron’s waveform characteristics, inter-spike-interval (ISI) distribution, and firing-rate distribution (Berke, 2008; Berke et al., 2004). First, we excluded putative neurons with waveforms that did have the pattern of large initial negative deflection (“valley”) followed by a smaller positivity (“peak”) that is typical of extracellular action potentials. Second, we required putative MSN waveforms to have a valley width > 100 μs, peak width > 350 μs, and average firing rate < 20 spikes per second. Third, we excluded neurons with an initial gap in their ISI distributions, which is typical of striatal “tonically active neurons” (TANs). Following these criteria, of 483 clearly isolatable waveforms, we categorized 281 as MSNs (171 from monkey P and 110 from monkey R) which were used for all data analyses.

Because we used acute recording techniques, in which the relative positions of neural tissue and recording contacts may drift over the course of a recording session, we do not claim that the CDh neurons recorded before and during inactivation are the same. Accordingly, we did not quantitatively compare the responses of individual neurons before on during inactivation, and instead compared population-level responses. In figure 2, we present example neuron data recorded before inactivation and qualitatively compared this to data recorded on the same contact during inactivation.

### Data analysis

All data analyses were performed in MATLAB (MathWorks, Natick, MA).

To minimize the impact of any slow fluctuations over the course of a recording session (Bondy et al., 2018), each neuron’s spike count data was z-scored separately in each non-overlapping successive “block-pair”. For example, blocks 1 and 2, comprising 136 (2 x 68) trials was considered a block-pair and included 68 trials with the cue contralateral to recorded CDh neurons (cue-contra) and 68 trials with cue-ipsi. To keep block-pairs non-overlapping, blocks 3 and 4 made up the next block-pair (each block contributed to only one block-pair); block-pairs also exclusively consisted of before inactivation or during inactivation data, never both. Separately for each neuron and for each of several window-durations, the mean and standard deviation of spike-count (or spike-rate) values was estimated in each block-pair; z-score was computed from spike-count/rate by subtracting the mean and dividing by the standard deviation. Z-scored spike counts were used in computing cue-side preferences and for classifier analyses.

To specifically quantify the effects of SC inactivation on cue-related modulation, we focused our analyses of neuronal activity on times before the motion-direction change. We previously found that many caudate MSNs display activity related to joystick-release or stimulus-change-related activity modulated by joystick-release (Arcizet and Krauzlis, 2018). Because SC inactivation systematically altered the probability of reporting the motion-direction change, alterations in CDh activity after the stimulus change could be secondary effects caused by changes in motor behavior, so these later epochs were excluded from analysis.

Logistic regression was used to separately test the effects of same-side and opposite-side SC inactivation on the proportion of significant single-neuron cue-side preferences. These analyses included cue-side (cue-contra or cue-ipsi), task-epoch (1-5), and inactivation state (before or during) as categorical predictors, giving each regression model 20 observations and 12 error degrees of freedom. Importantly, the regression model of same-side SC inactivation data resulted in a significant improvement over a constant model (χ^2^ vs constant model = 109, p = 1.19 × 10^−20^), but opposite-side did not (χ^2^ vs constant model = 9.24, p = 0.236).

Bootstrap tests were used to compare the distributions of ROC area values before and during SC inactivation. We computed the mean and skewness of ROC area distributions, then computed 95% confidence intervals by resampling with replacement 10000 times from measured ROC values to build distributions of mean and skewness and then calculating the 2.5^th^ and 97.5^th^ percentiles of those distributions; this was done separately on same-side and opposite-side inactivation data, and separately on before and during data. A significant difference in mean or skewness (noted as p < 0.05) was determined by finding non-overlapping confidence intervals. We note the effects of same-side SC inactivation on mean and skewness of ROC distributions in the results section; opposite-side SC inactivation caused a small but significant increase in mean (before: μ = 0.494, 95% CI = [0.488, 0.499]; during: μ = 0.506, 95% CI = [0.500, 0.512]; p < 0.05), and a nonsignificant change in skewness from negative to positive (before: skewness = −0.12, 95% CI = [-1.08, 0.42]; during: skewness = 0.12, 95% CI = [-0.17, 0.44]; p > 0.05).

Pearson correlation values were computed from trial-by-trial, z-scored spike counts between all pairs of simultaneously recorded neurons. Sessions with same-side SC inactivation plus recording yielded n_before_ = 4280 and n_during_ = 4042 pairs; opposite-side sessions yielded n_before_ = 2302 and n_during_ = 1468 pairs. Pairwise correlations were computed in each task epoch, separately for cue-contra and cue-ipsi conditions, and separately for before and during SC inactivation data. To quantify any variation in distributions of correlation values as a function of epoch, cue-side, inactivation state (before / during), or their interactions, we simultaneously fit distributions in each combination of conditions using the Stable family of distributions (Mandelbrot, 1960). This distribution has 4 parameters that govern its shape (α, β, c, and μ), which allows variation in the distribution’s central-tendency, width, skewness and the sharpness of its peak. To fit, we identified a single maximum likelihood solution **b** = [b_α_ b_β_ b_c_ b_μ_] to the equations: α = Xb_α_, β = Xb_β_, c = Xb_c_, and μ = Xb_μ_, where X is a categorical predictor array of experimental conditions including a constant term, main effects (epoch, cue-side, inactivation state), 2-way interactions, and 3-way interactions. We identified this solution (**b**) by minimizing the cost function: C = Σ-log(p(y| α, β, c, μ)) where y is a vector of all correlation values, and p is the Stable distribution PDF (probability density function). We computed **b** separately for correlations from same-side and opposite-side SC inactivations, and determined statistical significance by bootstrapping on **b** (we built distributions by shuffling and refitting 10000 times).

To statistically test the effects of inactivation on CDh neuron firing rates within and across sessions, we fit inter-spike interval (ISI) distributions with a generalized linear model (GLM) using inactivation state (before / during), session ID (1-13) and their interaction terms as categorical predictors. We assumed that ISIs were gamma distributed, and accordingly used a (canonical) negative inverse link function. We fit separate GLMs to same-side and opposite-side data, and both explained significantly more variance than a constant model (p ≪ 0.01). The same-side GLM included 476558 observations with 476540 error degrees of freedom, and opposite-side included 200168 observations with 200160 error degrees of freedom. In the results section, we report p-values from the main effect of inactivation state as indicating the significance of an across-sessions effect, and p-values from the interaction of inactivation state and individual session ID predictors as indicating the significance of a within-session effect.

To determine whether SC inactivation affected CDh population-level encoding of cue-side and task-epoch, we used a multidimensional classifier approach. We treated CDh neuronal data before and during SC inactivation separately, resulting in 26 datasets (before / during × 13 sessions), and always computed 5-fold cross-validated classifier performance (Hastie et al., 2013). We examined the performance of several types of classifier across datasets: (1) Linear discriminant, (2) Support Vector Machine (SVM), (3) naïve bayes, (4) decision tree, (5) k nearest neighbor (knn), (6) boosted decision tree, (7) boosted knn; each time a classifier was trained and tested on a dataset we used a Bayesian hyperparameter optimization procedure (Snoek et al., 2012). We selected the boosted decision tree classifier, using the AdaBoost method (Freund and Schapire, 1997), because we found it had the best performance across datasets.

## Acknowledgements

We thank B. Averbeck, B. Cumming, S. Goldstein, L. Katz, K. McAlonan, C. Quaia, L. Wang, and G. Yu for helpful discussions. This work was supported by the National Eye Institute Intramural Research Program at the National Institutes of Health.

## Supporting Material

**Fig. S1.**
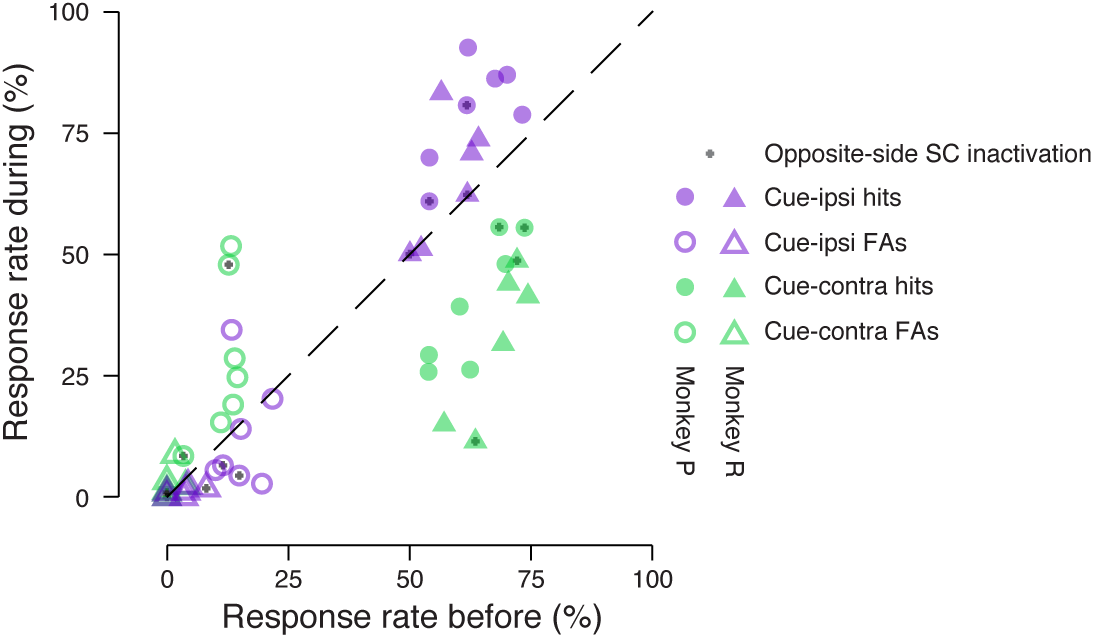
Performance summary comparing before and during SC inactivation broken down by SC inactivation side and subject. Hit (filled symbols) and false-alarm rates (empty symbols) before SC inactivation are plotted against rates from the same sessions during SC inactivation. Rates for cue-ipsilateral to SC inactivation site are plotted in purple and those for cue-contralateral are plotted in green. Triangles indicate rates for monkey R and circles indicate rates for monkey P. An added gray symbol plotted in the middle of rates distinguishes opposite-side SC inactivation sessions from same-side SC inactivation sessions.

## References

Akhlaghpour H, Wiskerke J, Choi JY, Taliaferro JP, Au J, Witten IB. 2016. Dissociated sequential activity and stimulus encoding in the dorsomedial striatum during spatial working memory. Elife 5:357. doi:10.7554/eLife.19507

Alexander GE, DeLong MR, Strick PL. 1986. Parallel organization of functionally segregated circuits linking basal ganglia and cortex. Annu Rev Neurosci 9:357–381. doi:10.1146/annurev.ne.09.030186.002041

Amita H, Hikosaka O. 2019. Indirect pathway from caudate tail mediates rejection of bad objects in periphery. Science Advances 5:eaaw9297. doi:10.1126/sciadv.aaw9297

Aosaki T, Graybiel AM, Kimura M. 1994a. Effect of the nigrostriatal dopamine system on acquired neural responses in the striatum of behaving monkeys. Science 265:412–415.

Aosaki T, Tsubokawa H, Ishida A, Watanabe K, Graybiel AM, Kimura M. 1994b. Responses of tonically active neurons in the primate’s striatum undergo systematic changes during behavioral sensorimotor conditioning. Journal of Neuroscience 14:3969–3984. doi:10.1523/JNEUROS-CI.14-06-03969.1994

Arcizet F, Krauzlis RJ. 2018. Covert spatial selection in primate basal ganglia. PLoS Biol 16:e2005930–28. doi:10.1371/journal.pbio.2005930

Averbeck BB, Lee D. 2006. Effects of noise correlations on information encoding and decoding. J Neurophysiol 95:3633–3644. doi:10.1152/jn.00919.2005

Bartolo-Orozco R, Saunders RC, Mitz A, Averbeck BB. 2019. Dimensionality, information and learning in prefrontal cortex. bioRxiv 823377.

Bisley JW, Goldberg ME. 2010. Attention, Intention, and Priority in the Parietal Lobe. Annu Rev Neurosci 33:1–21. doi:10.1146/annurev-neuro-060909-152823

Bogadhi AR, Bollimunta A, Leopold DA, Krauzlis RJ. 2019. Spatial Attention Deficits Are Causally Linked to an Area in Macaque Temporal Cortex. Current Biology 1–16. doi:10.1016/j.cub.2019.01.028

Bollimunta A, Bogadhi AR, Krauzlis RJ. 2018. Comparing frontal eye field and superior colliculus contributions to covert spatial attention. Nature Communications 9:3553. doi:10.1038/s41467-018-06042-2

Bondy AG, Haefner RM, Cumming BG. 2018. Feedback determines the structure of correlated variability in primary visual cortex. Nat Neurosci 79:1–15. doi:10.1038/s41593-018-0089-1

Bronfeld M, Belelovsky K, Erez Y, Bugaysen J, Korngreen A, Bar-Gad I. 2010. Bicuculline-induced chorea manifests in focal rather than globalized abnormalities in the activation of the external and internal globus pallidus. J Neurophysiol 104:3261–3275. doi:10.1152/jn.00093.2010

Carrasco M. 2011. Visual attention: The past 25 years. Vision Res 51:1484–1525. doi:10.1016/j.visres.2011.04.012

Cavanaugh J, Wurtz RH. 2004. Subcortical modulation of attention counters change blindness. J Neurosci 24:11236–11243. doi:10.1523/JNEUROSCI.3724-04.2004

Cohen MR, Kohn A. 2011. Measuring and interpreting neuronal correlations. Nat Neuro 14:811–819. doi:10.1038/nn.2842

Cohen MR, Maunsell JHR. 2009. Attention improves performance primarily by reducing interneuronal correlations. Nat Neurosci 12:1594–1600. doi:10.1038/nn.2439

Daw ND, Courville AC, Tourtezky DS, Touretzky DS. 2006. Representation and timing in theories of the dopamine system. Neural Comput 18:1637–1677. doi:10.1162/neco.2006.18.7.1637

Ding JB, Guzman JN, Peterson JD, Goldberg JA, Surmeier DJ. 2010. Thalamic Gating of Corticostriatal Signaling by Cholinergic Interneurons. Neuron 67:294–307. doi:10.1016/j.neuron.2010.06.017

Ding L, Gold JI. 2012. Separate, Causal Roles of the Caudate in Saccadic Choice and Execution in a Perceptual Decision Task. Neuron 75:865–874. doi:10.1016/j.neuron.2012.07.021

Ding L, Gold JI. 2010. Caudate Encodes Multiple Computations for Perceptual Decisions. Journal of Neuroscience 30:15747–15759. doi:10.1523/JNEUROSCI.2894-10.2010

Ecker AS, Berens P, Keliris GA, Bethge M, Logothetis NK, Tolias AS. 2010. Decorrelated neuronal firing in cortical microcircuits. Science 327:584–587. doi:10.1126/science.1179867

Freund Y, Schapire RE. 1997. A Decision-Theoretic Generalization of On-Line Learning and an Application to Boosting. Journal of Computer and System Sciences 55:119–139. doi:10.1006/jcss.1997.1504

Gershman SJ, Uchida N. 2019. Believing in dopamine. Nat Rev Neurosci 20:703–714. doi:10.1038/s41583-019-0220-7

Grunwerg BS, Krauthamer GM. 1992. Sensory responses of intralaminar thalamic neurons activated by the superior colliculus. Exp Brain Res 88:541–550. doi:10.1007/bf00228183

Hastie T, Tibshirani R, Friedman J. 2013. The Elements of Statistical Learning. New York, NY: Springer Science & Business Media. doi:10.1007/978-0-387-21606-5

Herman JP, Katz LN, Krauzlis RJ. 2018. Midbrain activity can explain perceptual decisions during an attention task. Nat Neuro 21:1651–1655. doi:10.1038/s41593-018-0271-5

Hikosaka O, Kim HF, Yasuda M, Yamamoto S. 2014. Basal ganglia circuits for reward value-guided behavior. Annu Rev Neurosci 37:289–306. doi:10.1146/annurev-neuro-071013-013924

Hikosaka O, Nakamura K, Nakahara H. 2006. Basal ganglia orient eyes to reward. J Neurophysiol 95:567–584. doi:10.1152/jn.00458.2005

Hikosaka O, Sakamoto M, Usui S. 1989. Functional properties of monkey caudate neurons. III. Activities related to expectation of target and reward. J Neurophysiol 61:814–832.

Jog MS, Kubota Y, Connolly CI, Hillegaart V, Graybiel AM. 1999. Building neural representations of habits. Science 286:1745–1749. doi:10.1126/science.286.5445.1745

Karnath H-O, Rorden C. 2012. The anatomy of spatial neglect. Neuropsychologia 50:1010–1017. doi:10.1016/j.neuropsychologia.2011.06.027

Kermadi I, Joseph JP. 1995. Activity in the caudate nucleus of monkey during spatial sequencing. J Neurophysiol 74:911–933. doi:10.1152/jn.1995.74.3.911

Kim HF, Hikosaka O. 2013. Distinct basal ganglia circuits controlling behaviors guided by flexible and stable values. Neuron 79:1001–1010. doi:10.1016/j.neuron.2013.06.044

Krauzlis RJ, Bogadhi AR, Herman JP, Bollimunta A. 2018. Selective attention without a neocortex. Cortex 102:161–175. doi:10.1016/j.cortex.2017.08.026

Krout KE, Loewy AD, Westby GW, Redgrave P. 2001. Superior colliculus projections to midline and intralaminar thalamic nuclei of the rat. J Comp Neurol 431:198–216. doi:10.1002/1096-9861(20010305)431:2<198::aid-cne1065>3.0.co;2-8

la Rocha de J, Doiron B, Shea-Brown E, Josic K, Reyes A. 2007. Correlation between neural spike trains increases with firing rate. Nature 448:802–806. doi:10.1038/nature06028

Lau B, Glimcher PW. 2008. Value representations in the primate striatum during matching behavior. Neuron 58:451–463. doi:10.1016/j.neuron.2008.02.021

Leavitt ML, Pieper F, Sachs AJ, Martinez-Trujillo JC. 2017. Correlated variability modifies working memory fidelity in primate prefrontal neuronal ensembles. Proc Natl Acad Sci USA 114:E2494–E2503. doi:10.1073/pnas.1619949114

Lee E, Seo M, Dal Monte O, Averbeck BB. 2015. Injection of a dopamine type 2 receptor antagonist into the dorsal striatum disrupts choices driven by previous outcomes, but not perceptual inference. J Neurosci 35:6298–6306. doi:10.1523/JNEUROSCI.4561-14.2015

Lovejoy LP, Krauzlis RJ. 2017. Changes in perceptual sensitivity related to spatial cues depends on subcortical activity. Proc Natl Acad Sci USA 114:6122–6126. doi:10.1073/pnas.1609711114

Lovejoy LP, Krauzlis RJ. 2010. Inactivation of primate superior colliculus impairs covert selection of signals for perceptual judgments. Nat Neurosci 13:261–266. doi:10.1038/nn.2470

Matsumoto N, Minamimoto T, Graybiel AM, Kimura M. 2001. Neurons in the thalamic CM-Pf complex supply striatal neurons with information about behaviorally significant sensory events. J Neurophysiol 85:960–976. doi:10.1152/jn.2001.85.2.960

May PJ, McHaffie JG, Stanford TR, Jiang H, Costello MG, Coizet V, Hayes LM, Haber SN, Redgrave P. 2009. Tectonigral projections in the primate: a pathway for pre-attentive sensory input to midbrain dopaminergic neurons. Eur J Neurosci 29:575–587. doi:10.1111/j.1460-9568.2008.06596.x

McHaffie JG, Stanford TR, Stein BE, Coizet V, Redgrave P. 2005. Subcortical loops through the basal ganglia. Trends Neurosci 28:401–407. doi:10.1016/j.tins.2005.06.006

Minamimoto T, Kimura M. 2002. Participation of the thalamic CM-Pf complex in attentional orienting. J Neurophysiol 87:3090–3101. doi:10.1152/jn.2002.87.6.3090

Mink JW. 2001. Basal ganglia dysfunction in Tourette’s syndrome: a new hypothesis. Pediatr Neurol 25:190–198. doi:10.1016/s0887-8994(01)00262-4

Mitchell JF, Sundberg KA, Reynolds JH. 2009. Spatial attention decorrelates intrinsic activity fluctuations in macaque area V4. Neuron 63:879–888. doi:10.1016/j.neuron.2009.09.013

Miyachi S, Hikosaka O, Lu X. 2002. Differential activation of monkey striatal neurons in the early and late stages of procedural learning. Exp Brain Res 146:122–126. doi:10.1007/s00221-002-1213-7

Miyashita N, Hikosaka O, Kato M. 1995. Visual hemineglect induced by unilateral striatal dopamine deficiency in monkeys. NeuroReport 6:1257–1260.

Moore T, Zirnsak M. 2017. Neural Mechanisms of Selective Visual Attention. Annu Rev Psychol 68:47–72. doi:10.1146/annurev-psych-122414-033400

Moreno-Bote R, Beck J, Kanitscheider I, Pitkow X, Latham PE, Pouget A. 2014. Information-limiting correlations. Nat Neurosci 17:1410–1417. doi:10.1038/nn.3807

Müller JR, Philiastides MG, Newsome WT. 2005. Microstimulation of the superior colliculus focuses attention without moving the eyes. Proc Natl Acad Sci USA 102:524–529. doi:10.1073/pnas.0408311101

Nakajima M, Schmitt LI, Halassa MM. 2019. Prefrontal Cortex Regulates Sensory Filtering through a Basal Ganglia-to-Thalamus Pathway. Neuron 103:445–458.e10. doi:10.1016/j.neuron.2019.05.026

Nakano K, Kayahara T, Tsutsumi T, Ushiro H. 2000. Neural circuits and functional organization of the striatum. Journal of Neurology 247 Suppl 5:V1–15. doi:10.1007/pl00007778

Nelson AB, Kreitzer AC. 2014. Reassessing Models of Basal Ganglia Function and Dysfunction. Annu Rev Neurosci 37:117–135. doi:10.1146/annurev-neuro-071013-013916

Nevet A, Morris G, Saban G, Arkadir D, Bergman H. 2007. Lack of spike-count and spike-time correlations in the substantia nigra reticulata despite overlap of neural responses. J Neurophysiol 98:2232–2243. doi:10.1152/jn.00190.2007

Nienborg H, Cohen MR, Cumming BG. 2012. Decision-related activity in sensory neurons: correlations among neurons and with behavior. Annu Rev Neurosci 35:463–483. doi:10.1146/annurev-neuro-062111-150403

Pachitariu M, Steinmetz N, Kadir S, Carandini M, Harris KD. 2016. Kilosort: realtime spike-sorting for extracellular electrophysiology with hundreds of channels. bioRxiv. doi:10.1101/061481

Rao RPN. 2010. Decision making under uncertainty: a neural model based on partially observable markov decision processes. Front Comput Neurosci 4:146. doi:10.3389/fncom.2010.00146

Ravel S, Legallet E, Apicella P. 1999. Tonically active neurons in the monkey striatum do not preferentially respond to appetitive stimuli. Exp Brain Res 128:531–534.

Redgrave P, Coizet V, Comoli E, McHaffie JG, Leriche M, Vautrelle N, Hayes LM, Overton P. 2010. Interactions between the Midbrain Superior Colliculus and the Basal Ganglia. Front Neuroanat 4. doi:10.3389/fnana.2010.00132

Reynolds JH, Chelazzi L. 2004. Attentional Modulation of Visual Processing. Annu Rev Neurosci 27:611–647. doi:10.1146/annurev.neuro.26.041002.131039

Saint-Cyr JA, Ungerleider LG, Desimone R. 1990. Organization of visual cortical inputs to the striatum and subsequent outputs to the pallido-nigral complex in the monkey. J Comp Neurol 298:129–156. doi:10.1002/cne.902980202

Samejima K, Ueda Y, Doya K, Kimura M. 2005. Representation of action-specific reward values in the striatum. Science 310:1337–1340. doi:10.1126/science.1115270

Seo M, Lee E, Averbeck BB. 2012. Action Selection and Action Value in Frontal-Striatal Circuits. Neuron 74:947–960. doi:10.1016/j.neuron.2012.03.037

Shadlen MN, Britten KH, Newsome WT, Movshon JA. 1996. A computational analysis of the relationship between neuronal and behavioral responses to visual motion. Journal of Neuroscience 16:1486–1510.

Sheng M-J, Lu D, Shen Z-M, Poo M-M. 2019. Emergence of stable striatal D1R and D2R neuronal ensembles with distinct firing sequence during motor learning. Proc Natl Acad Sci USA 116:11038–11047. doi:10.1073/pnas.1901712116

Snoek J, Larochelle H, Adams RP. 2012. Practical Bayesian Optimization of Machine Learning Algorithms In: Pereira F, Burges CJC, Bottou L, Weinberger KQ, editors. Advances in Neural Information Processing Systems 25. Curran Associates, Inc. pp. 2951–2959.

Treue S. 2001. Neural correlates of attention in primate visual cortex. Trends Neurosci 24:295–300. doi:10.1016/s0166-2236(00)01814-2

White JK, Monosov IE. 2016. Neurons in the primate dorsal striatum signal the uncertainty of object-reward associations. Nature Communications 7:12735. doi:10.1038/ncomms12735

Yamamoto S, Kim HF, Hikosaka O. 2013. Reward value-contingent changes of visual responses in the primate caudate tail associated with a visuomotor skill. J Neurosci 33:11227–11238. doi:10.1523/JNEUROSCI.0318-13.2013

Yasuda M, Hikosaka O. 2015. Functional territories in primate substantia nigra pars reticulata separately signaling stable and flexible values. J Neurophysiol 113:1681–1696. doi:10.1152/jn.00674.2014

Zénon A, Krauzlis RJ. 2012. Attention deficits without cortical neuronal deficits. Nature 489:434–437. doi:10.1038/nature11497

